# Structural survey of HIV-1-neutralizing antibodies targeting Env trimer delineates epitope categories and suggests vaccine templates

**DOI:** 10.1101/312579

**Authors:** Gwo-Yu Chuang, Jing Zhou, Reda Rawi, Chen-Hsiang Shen, Zizhang Sheng, Anthony P. West, Baoshan Zhang, Tongqing Zhou, Robert T. Bailer, Nicole A. Doria-Rose, Mark K. Louder, Krisha McKee, John R. Mascola, Pamela J. Bjorkman, Lawrence Shapiro, Peter D. Kwong

## Abstract

HIV-1 broadly neutralizing antibodies are desired for their therapeutic potential and as templates for vaccine design. Such antibodies target the HIV-1-envelope (Env) trimer, which is shielded from immune recognition by extraordinary glycosylation and sequence variability. Recognition by broadly neutralizing antibodies thus provides insight into how antibody can bypass these immune-evasion mechanisms. Remarkably, antibodies neutralizing >25% of HIV-1 strains have now been identified that recognize all major exposed surfaces of the prefusion-closed Env trimer. Here we analyzed all 206 broadly neutralizing antibody-HIV-1 Env complexes in the PDB with resolution suitable to define their interaction chemistries. These segregated into 20 antibody classes based on ontogeny and recognition, and into 6 epitope categories (*V1V2, glycan-V3, CD4-binding site, silent face center, fusion peptide*, and *subunit interface*) based on recognized Env residues. We measured antibody neutralization on a 208-isolate panel and analyzed features of paratope and B cell ontogeny. The number of protruding loops, CDR H3 length, and level of somatic hypermutation for broadly HIV-1 neutralizing antibodies were significantly higher than for a comparison set of non-HIV-1 antibodies. For epitope, the number of independent sequence segments was higher (P < 0.0001), as well as the glycan component surface area (P = 0.0005). Based on B cell ontogeny, paratope, and breadth, the CD4-binding site antibody IOMA appeared to be a promising candidate for lineage-based vaccine design. In terms of epitope-based vaccine design, antibody VRC34.01 had few epitope segments, low epitope-glycan content, and high epitope-conformational variability, which may explain why VRC34.01-based design is yielding promising vaccine results.

## Significance

Over the last decade, structures of broadly neutralizing antibodies have been determined, encompassing all major exposed surfaces on the prefusion-closed HIV-1-envelope (Env) trimer. To provide insight into how the immune system generated these antibodies along with molecular detail of their recognition, we surveyed known Env-antibody structures. We found that (i) broadly neutralizing antibodies recognized sequence variability and glycan at frequencies similar to their average occurrence on the Env-trimer surface, (ii) almost two-thirds of the paratopes utilized protruding loops, which were otherwise less common; and (iii) the broadest HIV-1-neutralizing antibodies had high levels of SHM, and the most potent unusual recombination, but antibodies with more reproducible B cell ontogenies still neutralized broadly, and these may be better templates for re-elicitation.

Antibodies against HIV-1 that neutralize a significant fraction of the diverse primary isolates that typify HIV-1 transmission are highly sought. These antibodies target the HIV-1 envelope (Env) trimer, which is comprised of gp120 and gp41 subunits and shielded from immune recognition by extraordinary glycosylation (1), sequence variability (2), and conformational masking (3). A decade ago, only four such broadly neutralizing antibodies had been identified. However, in 2009, the identification by B cell culture of antibody PG9 (along with a somatic variant PG16) (4) and in 2010 the identification by probe-based sorting of antibody VRC01 (along with somatic variants VRC02 and VRC03) (5) initiated an outbreak of discovery, with dozens of broadly neutralizing antibodies identified by culture or probe-based sorting of B cells from HIV-1 infected donors (reviewed in (6)).

Importantly, structures for many of these antibodies in complex with HIV-1 Env have been determined, which link antibody function (breadth and potency of neutralization) with molecular features of antibody recognition (paratope) and chemical details of Env-recognized site (epitope). For example, structures of PG9 with scaffolded V1V2 regions of Env (7) and also with Env trimer (8) have revealed a protruding anionic tyrosine-sulfated loop penetrating the glycan shield to interact with a conserved cationic site on V2; similarly structures of VRC01 with core gp120 (9) or with a fully glycosylated Env trimer (10) have revealed how SHM-based optimization of interactions – especially those involving glycan – allows for broad recognition of the CD4-binding site. Most recently, the structure of antibody VRC-PG05 in complex with a glycosylated Env core (11) has shown that even the highly glycosylated center of the Env “silent face” could be recognized by broadly neutralizing antibodies.

The structure of VRC-PG05 in complex with Env completes recognition by structurally characterized antibody of all major exposed surfaces on the prefusion-closed HIV-Env trimer.

Here we take advantage of this completion to perform a structure-based survey of all antibody-HIV-1 Env structures in the Protein Data Bank (PDB). We limited analyses to structures of human antibodies (with neutralization breadth > 25%) in complex with epitopes on the prefusion-closed Env trimer with resolutions sufficient to define side chain orientation and interactions. The focus on the prefusion-closed conformation of Env alleviated issues of conformational masking and enabled comparisons on a single structural entity. We further classified structures by antibody class (B cell ontogeny and structural mode of recognition) to unique classes and used a bottom-up classification based on residues of each epitope to define epitope categories. We measured antibody neutralization on a diverse cross-clade panel of 208 HIV-1 isolates and correlated structural features of epitope and paratope with functional characteristics of neutralization breadth and potency. We also analyzed paratopes for ease of lineage re-elicitation and epitopes for ease of mimicry. Overall, our structural survey of HIV-1 antibodies targeting the prefusion-closed Env trimer delineates categories, identifies underlying relationships between structural properties of paratope and epitope and functional properties of neutralization, and suggests favorable candidates for vaccine design.

## Results

### Broadly neutralizing antibodies that recognize the prefusion-closed Env trimer segregate into 20 classes and 6 categories

The PDB contains over 200 coordinate sets for human antibodies in complex with various portions of HIV-1 Env, ranging from fully glycosylated Env trimers to core gp120, scaffolded domains, representative *N*-linked glycans, and peptide fragments (**Dataset S1**). We chose to limit our analysis to structures with resolutions sufficient to define chemistry of interaction: for X-ray crystal structures, we used a resolution limit of 3.9 Å, and for cryo-electron microscopy structures, we used a resolution limit of 4.5 Å. Also, we chose to limit our analysis to antibodies that neutralize at least 25% of HIV-1 at 50 μg/ml, as measured in a cross-clade panel of at least 100 isolates. Finally, to alleviate issues of conformation, we limited our analysis to only those antibodies that recognize the prefusion-closed conformation of Env as this allowed us to focus on a single structural entity: the Env trimer in its prefusion-closed state.

We next used similarity in B cell ontogeny and structural mode of recognition to classify these PDBs into antibody classes (12) (**Table 1**). While some antibody classes have only been found in a single donor to date (e.g., CH103, VRC38), other classes have been found in more than one donor (e.g. VRC01, 8ANC131). The largest class involved the VRC01 class (13), with 53 PDBs describing 33 antibodies. Antibodies recognizing glycan-V3, derived primarily from classes that have only a single known clonal lineage, although PGT121 and BG18 lineages do share the same D3-3 gene and mode of recognition (14, 15). For antibodies recognizing the spike apex, prior analyses placed antibody lineages PG9, PGT145 and CH01 in the same class (7). While structural information showed parallel strand recognition for PG9 and CH03 indicating similar classes (16), structures of PGT145 with Env trimer (17, 18) indicated loop insertion into a trimeric hole at the spike apex, and hence a distinct mode of recognition and class for PGT145. We used the name of the first identified antibody in each antibody class as the name of the class.

**Table 1.**
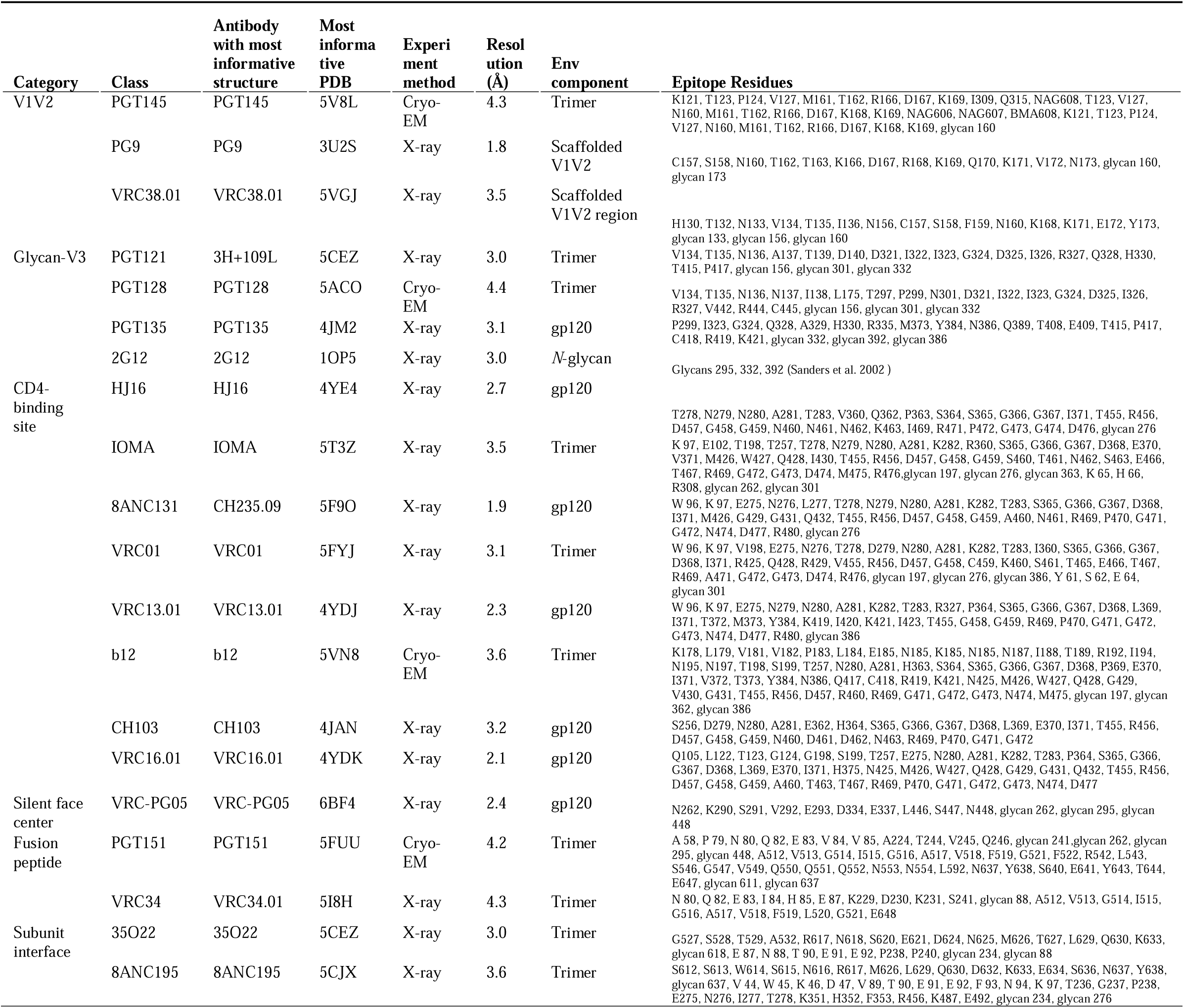
Structural details for 20 classes of antibodies in complex with HIV-1 Env.

We selected a structural representative antibody for each class based on the most informative structure (for example, choosing an antibody-Env trimer complex over the same antibody with a deglycosylated core gp120) (**Fig. 1A**) Based on the 20 representative antibody-antigen complex structures, we defined epitope residues on the prefusion-closed Env trimer by buried surface area calculations (see methods). For 2G12, the most complete antibody-Env complexes defined interactions with only a few *N*-linked glycans (19); therefore, we used published biochemical studies to define the 2G12 epitope residues (20). Next, we carried out hierarchical clustering of epitope residues, which clustered the 20 antibody epitopes into 6 categories (**Fig. 1A,B**). Three of the categories had been previously described: V1V2, glycan-V3 and CD4-binding site, and each of these categories encompassed at least three different antibody classes. Three additional categories were identified, including the “silent face center” (with antibody VRC-PG05), “fusion peptide” (with antibodies PGT151 and VRC34), and “subunit interface” (with antibodies 35O22 and 8ANC195). Beyond shared epitope residues, each of the categories did appear to have specific characteristics. For example, smFRET measurements define three prefusion states (states 1-3) of the Env trimer and found broadly neutralizing antibodies to preferentially recognize state 1 (21); only two antibodies, VRC34.01 (22) and PGT151 (personal communications, Walther Mothes), have thus far been found to shift the smFRET population to state 2, and both were assigned to the same epitope category. Thus, our structural analysis defined six categories of broadly neutralizing antibodies, and antibodies in each category generally shared similar characteristics.

**Fig. 1.**
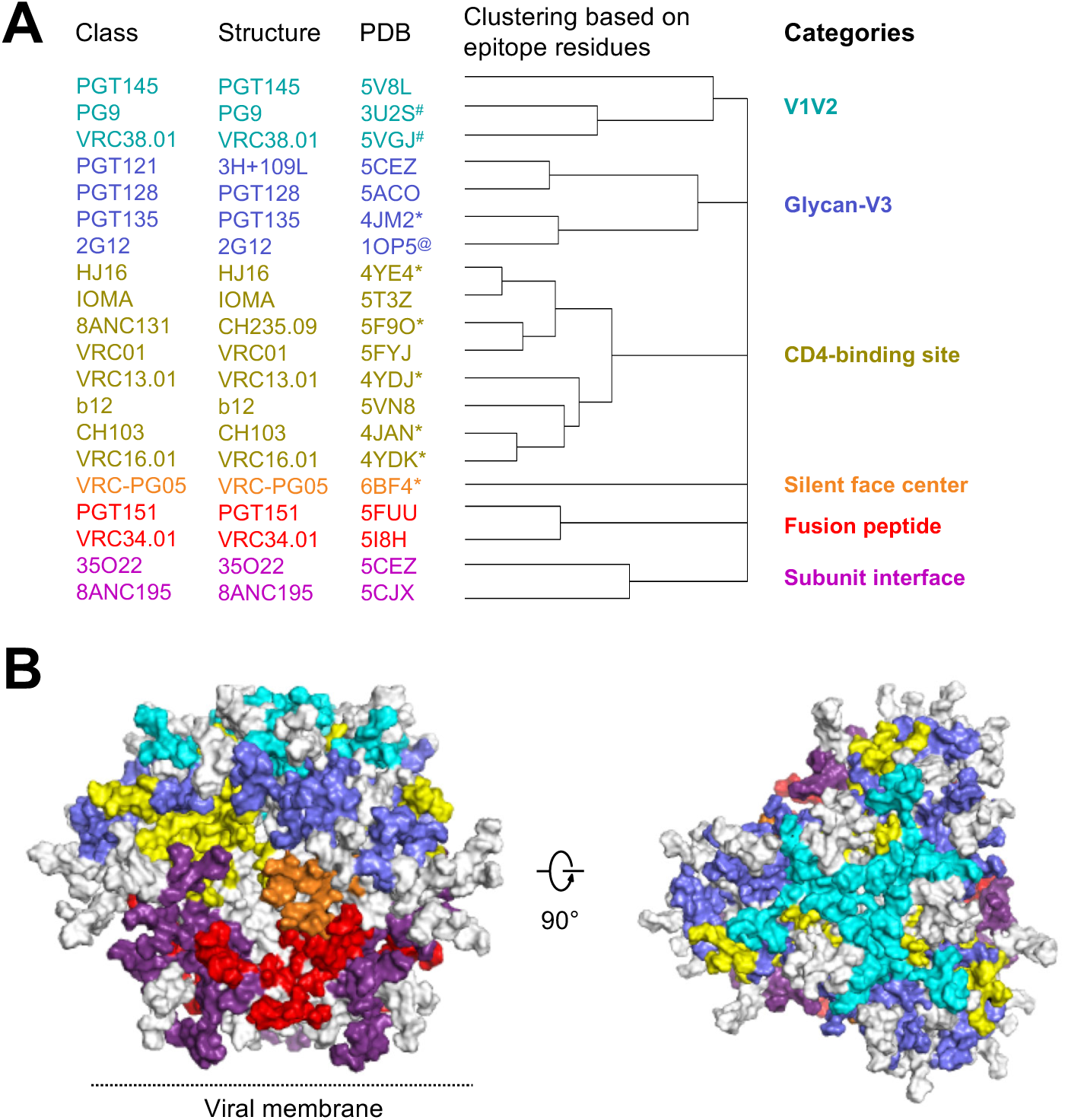
20 classes of broadly neutralizing antibodies recognize the prefusion-closed Env trimer and segregate into 6 categories based on the Env residues with which they interact. (**A**) All HIV-1 Env-antibody complexes structures in the PDB (Table S1) were assigned to classes (leftmost column, listed by the name of first reported antibody of the class) based on similarities in B cell ontogeny and mode of recognition, with a representative structure and PDB for the class (2nd and 3rd columns from left), which were chosen based on resolution and degree to which Env in the structure resembled prefusion-closed trimer; “*” indicate structures determined in gp120-core context; “#” indicates structures determined with V1/V2 scaffold; “@” indicates glycan N295, N332, and N392 used for 2G12 epitope (see methods) (**B**) Prefusion-closed Env trimer with molecular surface colored by categories defined in (A). Epitope residues shared by antibodies in separate classes were colored according to surface area and requirements for antibody binding. For example, glycan N276 has interactions with antibody 8ANC195 of the subunit interface category and with multiple antibodies of the CD4-binding site category; however, because glycan N276 is required for 8ANC195, and generally only accommodated by antibodies that target the CD4-binding site, we colored glycan N276 to be part of the subunit interface category. Left image is shown with viral membrane at bottom; right image is rotated 90° to look down on the trimer apex.

### Neutralization characteristics for 20 antibody classes

As the most important functional property of the 20 antibody classes is their ability to neutralize diverse HIV-1 isolates, we felt it important to measure accurately this feature in a way that allowed comparison between different antibodies. We thus measured both neutralization breadth and potency for the representative member of each class on a diverse cross-clade panel of 208 HIV-1 isolates (23) (**Fig. 2A,B, Data Set S2**). For functional representative, we chose the antibody-class member with highest neutralization breadth. For the PGT121 class, we chose antibody 10-1074 (14); for the VRC01 class, we chose antibody N6 (24); and for the 8ANC131 class we chose antibody CH235.12 (25). Notably, the newly measured median neutralization IC50 generally differed by less than 2 fold from prior reported values (5, 7, 11, 22, 24-40), with the breadth for all 20 classes averaging 56.7%, with an average geometric mean IC_50_ of 0.27 μg/ml.

**Fig. 2.**
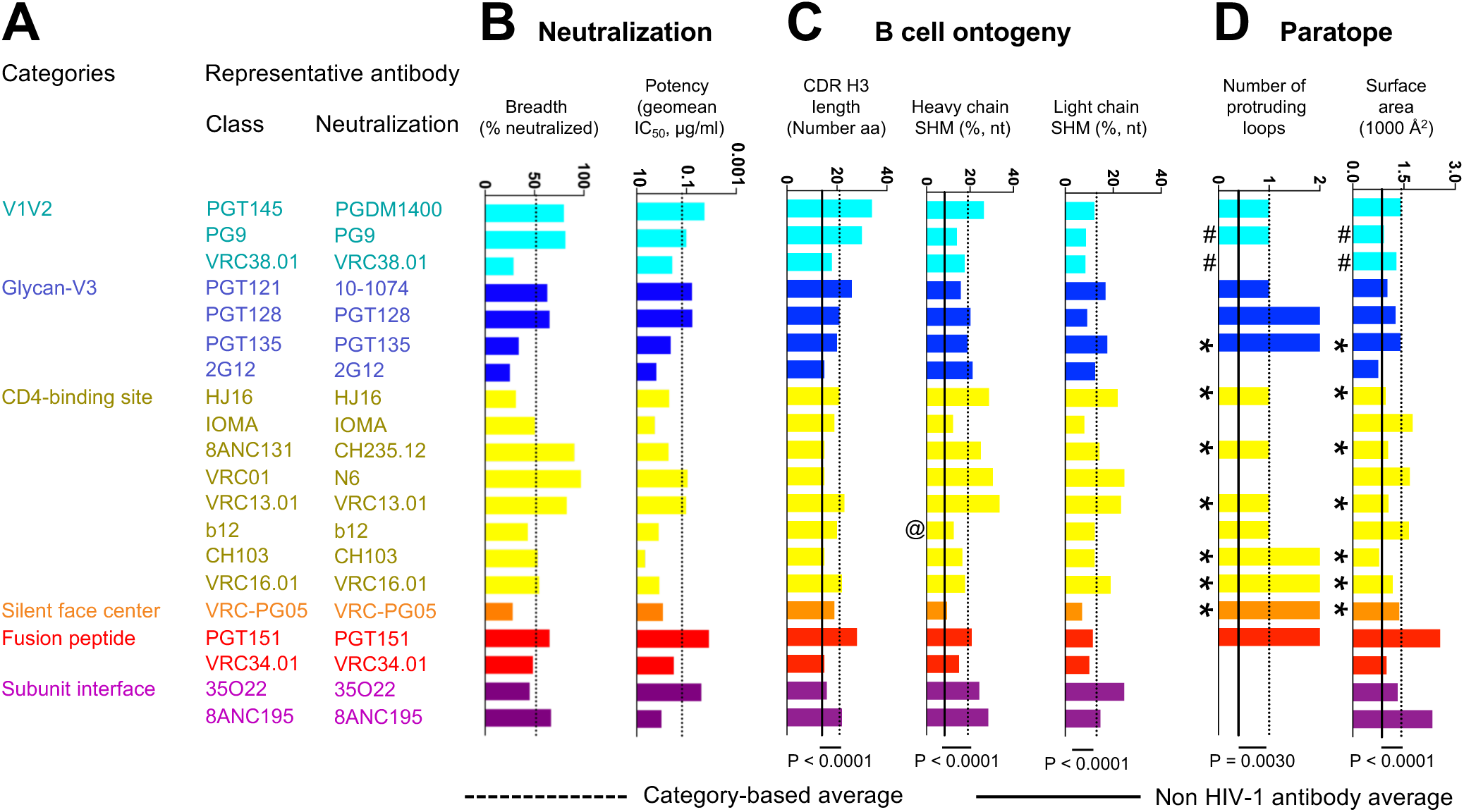
Neutralization, B cell ontogeny, and paratope characteristics for 20 classes of broadly neutralizing antibodies. **(A)** The broadest neutralizing antibody from each class was chosen as representative. **(B)** Neutralization properties. Bar graphs show breadth and potency of each representative antibody on the 208-isolate panel, with dotted line showing averages. **(C)** Features of B cell ontogeny. CDR H3 lengths are defined based on Kabat nomenclature. @ indicates that antibody b12 is derived from a phage library. IOMA is the only antibody with CDHR H3 length under 20, SHM <12%, and breadth approaching 50%, and thus may be a promising candidate for lineage-based vaccine design. By contrast, VRC01-class antibodies have high SHM and PG9/PGT145 have unusually long CDR H3, complicating their lineage-based elicitation. **(D)** Paratope properties. “*” indicate structures determined in gp120-core context; “#” indicates structures determined with V1/V2 scaffold. Structures denoted with “*” or “#” were included in calculations of the category averages for protruding loop numbers, but not for paratope surface area. P-values were determined using two-tailed Mann-Whitney test.

### B cell ontogenies, paratopes, and epitopes of antibodies recognizing the prefusion-closed Env trimer

B cell ontogenies involve the recombination events that lead to creation of the ancestor B cell for each lineage, as well as SHM that lead to mature antibody clones. To estimate elicitation likelihoods related to recombination and SHM, we extracted CDR H3 length and V-gene SHM (**Fig. 2C**). We observed that the average CDR H3 length and heavy chain SHM of the 20 HIV-1 broadly neutralizing antibodies were higher than those of next generation sequencing (NGS) reads from three healthy donors (P<0.0001) (25).

We also extracted critical features of the paratope (**Fig. 2D**). We selected a measure related to recognition mode (number of protruding antibody loops) and a second measure related to recognition extent (the surface area of the interacting paratope). Thirteen of the 20 antibodies utilized protruding loops in recognition, although this feature was less common in non-HIV-1 antibodies (as defined on a set of 81 non-HIV-1 antibody-antigen structures, see **Datasets S3** and **S4**). The paratope surface areas were almost twice as high as observed with non-HIV-1 antibodies (P<0.0001), and this may relate to glycan interactions, as antibodies with deglycosylated cores generally showed lower interactive surface areas.

For epitope, we measured several features including sequence conservation, extent of glycan contribution, number of independent sequence elements in each epitope, and the structural variability of the epitope (**Fig. 3A-D**). For sequence conservation, epitopes of broadly neutralizing antibodies (75% conservation) were similar to the average conservation of the closed trimer (69%). For glycan, we observed the average contribution to epitope surface area (45%) was similar to the surface area contribution of MAN5 glycans to the overall Env trimer surface area (53%). Also, we observed that the average epitope glycan composition (P=0.0005), total epitope surface area (P<0.0001), and number of epitope segments (P<0.0001) of the 20 HIV-1 broadly neutralizing antibodies were higher than those of non-HIV-1 antibodies (**Datasets S3** and **S4**; for glycan composition specifically, the comparison was done with a non-HIV-1 glycoprotein set with 16 antibody antigen complexes, see Methods and **Dataset 5**). It may be that features common in HIV-1 neutralizing antibodies, but generally rare, provide insight into recognition constraints: in this case suggesting that HIV broadly neutralizing antibodies were selected specifically to overcome sequence variability and glycosylation.

**Fig. 3.**
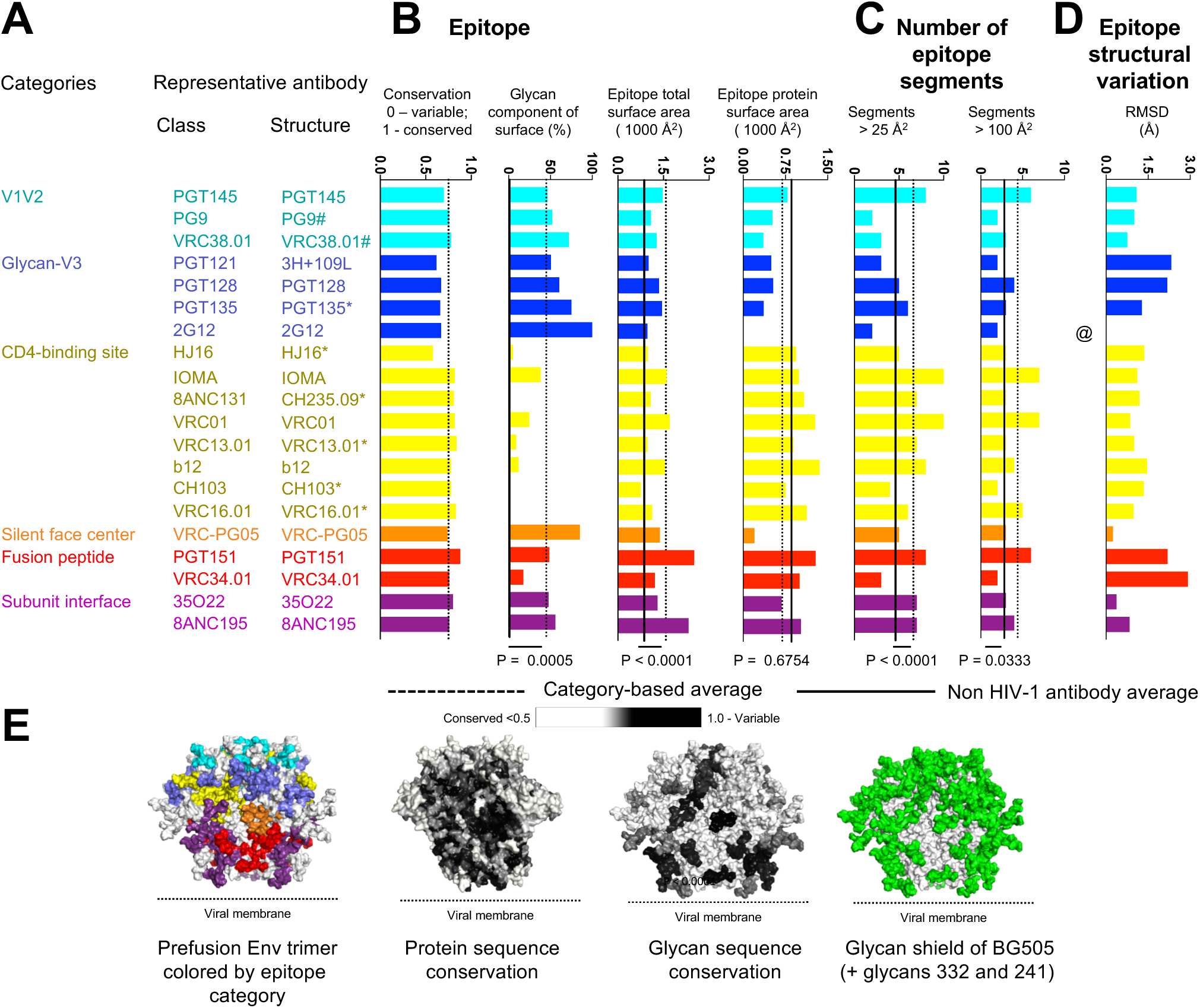
Epitope features for 20 classes of broadly neutralizing antibodies. **(A)** Categories, classes, and representative structures. “*” indicate structures determined in gp120-core context; “#” indicates structures determined with V1/V2 scaffold; @ indicates that the epitope structural variation of antibody 2G12 was not calculated due to the lack of protein epitope. Structures denoted with “*” or “#” were not included in calculations of the category averages for number of epitope segments and epitope total surface. Structures denoted with “*” were not included in calculations of the category average for epitope glycan composition. **(B)** Epitope properties. Bar graphs show conservation and glycan-interactive surface for each representative antibody, with dotted line showing averages. “*” indicate structures determined in gp120-core context, where glycan surface areas may not reflect trimer interactions. **(C)** Number of epitope segments, shown for segments with >25 Å2 or > 100 Å2 surface area. VRC34 has low number epitope segments and low glycan context (black arrow), and this may explain why fusion peptide immunization are yielding promising results from epitope-based vaccine design. (**D**) Observed variation in epitope structure, as calculated from RMSD of recognized amino acid residues, computed pairwise among class members of each category and unliganded trimer. (**E**) Prefusion-closed Env trimer colored by epitope and by various epitope features. P-values were determined using two-tailed Mann-Whitney test.

### Underlying relationships between neutralization, paratope and epitope

To provide insight into other features specific to these HIV-1 neutralizing antibodies, we correlated neutralization with epitope and paratope features to reveal underlying relationships (**Fig. 4A**). Although a number of features correlated with p-values of less than 0.05, when corrected for multiple comparison, few correlations were significant. Significant correlations included the expected positive correlation between surface area of paratope and surface area of epitope (P < 0.0001, r = 0.9845) (**Fig. 4B**) and a negative correlation between protein and glycan surface areas (P = 0.0002, r = −0.8413) (**Fig. 4C**). This negative correlation suggests antibodies to have an overall surface area that can comprise either protein or glycan; thus, a glycan-focused antibody generally has lower protein-interactive surface than a protein-focused antibody, and vice versa.

**Fig. 4.**
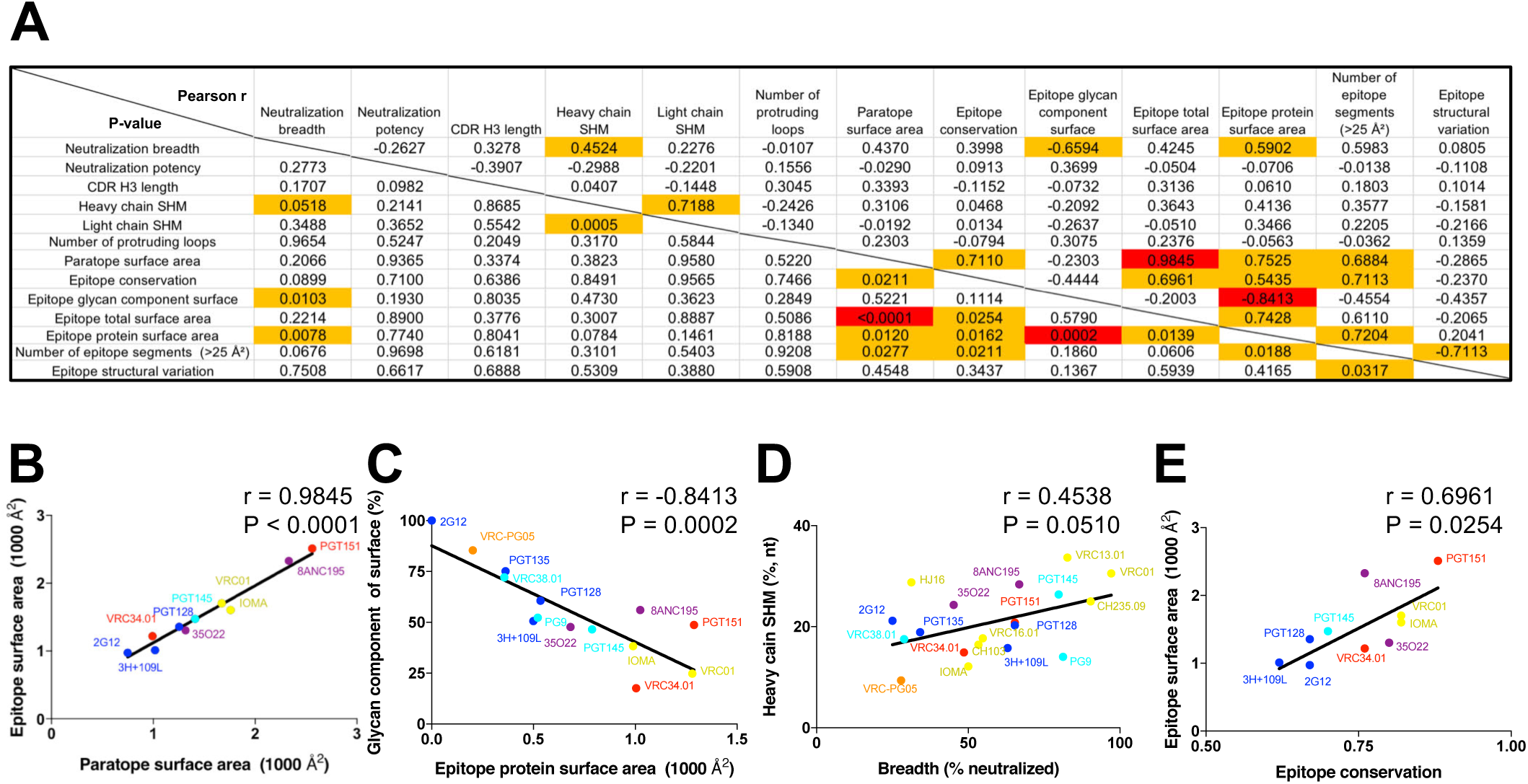
Correlations of neutralization and features of epitope and paratope reveal underlying relationships. Correlation matrices are shown for properties of neutralizing antibodies that recognize the prefusion closed Env trimer. (**A**) Pearson correlations (r) are provided in the upper right; P-values (not adjusted for multiple comparisons) are provided in the lower left. Correlations with P-value of less than 0.06 were highlighted with dark red font, and correlations with P-value of less than 0.05 after Bonferroni correction were highlighted with both dark red font and orange fill. (**B**) Correlation between paratope surface area and epitope surface area. (**C**) Correlation between Epitope protein surface area and glycan component of surface. (**D**) Correlation between neutralizing breadth and heavy chain SHM. (**E**) Correlation between epitope conservation and epitope total surface area.

Other correlations did not achieve statistical significance, but were potentially revealing. For example, we observed antibody neutralization breadth to correlate positively with degree of heavy chain SHM (P = 0.0510, r = 0.4538) and buried protein-epitope surface area (P = 0.0079, r = 0.5892) and negatively with degree of epitope glycan (P = 0.0105, r = −0.6579) (**Fig. 4D**), indicating that while antibody recognition can include glycan, this nevertheless reduces breadth. Meanwhile sequence conservation correlated with size of the epitope (P = 0.0254, r = 0.6961)/paratope (P = 0.0211, r = 0.7110) (**Fig. 4E**), suggesting that one way to overcome variation is by reducing the area of recognized surface. We also noticed that the broadest HIV-1-neutralizing antibodies had high levels of SHM and the most potent had unusual CDR H3s (**Fig. 2**). Overall, these observations suggested both the frequency and the extent that neutralizing antibodies targeting the prefusion-closed Env trimer utilize protruding loops, unusual recombination, and extensive SHM to overcome immune evasion mechanism of extraordinary glycosylation and high sequence variability.

### Re-elicitation: Antibody-lineage based vaccine design

We also analyzed antibodies to identify which B cell ontogenies might be most easily re-elicited by antibody-lineage based vaccine design. Such design is premised on the re-elicitation of antibodies with similar ontogenies, through priming of ancestor B cells and induction of their maturation (16, 41, 42). Two factors affect re-elicitation: the likelihood that an appropriate recombination event produces an appropriate unmutated common ancestor (UCA) and the likelihood that this UCA will mature through processes of SHM to achieve similar functional properties of neutralization. The former is affected by prevalence of appropriate V-genes and by the contribution to the paratope of CDR H3 features such as D-gene and N-nucleotide addition, which we calculated for each of the 20 classes of neutralizing antibodies (**Fig. 5**). We also calculated features of SHM, such as the percent of the V gene-contributed paratope surface altered by SHM or the prevalence of rare mutations (43). Although some of the broadest and most potent antibodies had features that lowered the probability of their re-elicitation, there were antibody classes with less rare SHM and CDR H3 characteristics such as IOMA, as well as antibodies like PG9 with extensive D-gene contributions to antigen recognition that might be re-elicited with greater ease.

**Fig. 5.**
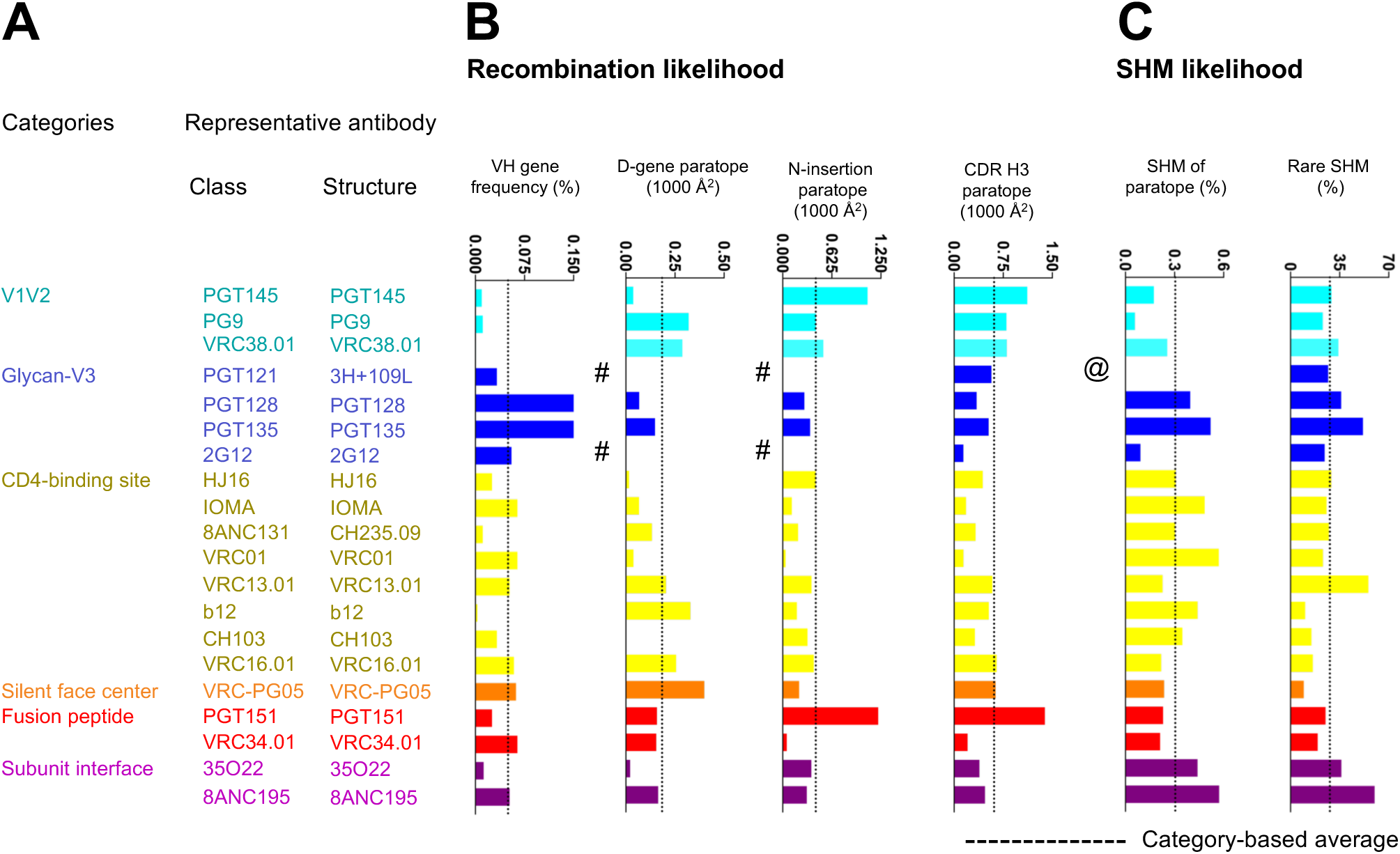
Factors affecting re-elicitation. **(A)** Categories, classes, and representative structures. **(B)** Recombination properties. Bar graphs show average VH gene frequency and paratope properties of D gene, N addition and CDR H3 for each representative antibody. Average VH gene frequencies report from healthy donors in PRJNA304166. D gene and N additions occurring at the junctions are from IMGT V-quest (http://www.imgt.org/IMGT_vquest/vquest). # indicates that antibody missing nucleotide sequences at junction. **(C)** SHM likelihood. Bar graphs show the paratope contributed from SHM (1st amino acid to the 2nd conserved Cys) and rare SHM. @ indicates that antibody missing nucleotide sequence.

## Discussion

The completion by antibody VRC-PG05 of structural characterization of the recognition by broadly neutralizing antibodies of all major exposed surfaces on the prefusion-closed Env trimer enables both comprehensive and aggregate-level analyses: Relative to paratope, we could address which characteristics allow for broad recognition of the glycosylated and sequence-variable Env trimer, and relative to the Env trimer surface, we could address which features were recognized and which were avoided. For paratope, our results indicated protruding loops and extensive SHM to dominate recognition (an observation previously reported by Klein and colleagues (44) utilizing a less complete set of HIV-1-directed antibodies), although antibodies with more common features could be found. For epitope, our results indicate both glycan and sequence variation to be recognized by antibody, which may relate to the extent that these features dominate the surface of the prefusion-closed Env trimer.

While our focus on the prefusion-closed trimer allowed us to relate properties of recognized epitopes to average properties of the Env trimer in a specific conformation, this focus led to the omission from our analysis of an important class of broadly neutralizing antibodies, those that target the membrane-proximal external region (MPER) (**Fig. S1** and **Table S2**). MPER antibodies generally show low levels of recognition of the native Env trimer (45) and have thus been proposed to recognize Env trimer in a different conformation (46). In addition to differences in conformation of Env recognized, MPER antibodies also target regions of Env with significantly lower glycan and sequence variability. Thus, the immune-evasion mechanisms that shield the MPER appear to differ from those shielding the prefusion-closed Env trimer, and the antibody mechanisms that allow for recognition of the MPER likely differ as well from those that allow recognition of the closed Env trimer.

In terms of antibody templates most favorable for vaccine design, our analysis suggested the IOMA lineage as being most favorable for re-elicitation by antibody lineage-based approaches due to the lack of unusual features in its B cell ontogeny. Furthermore, in comparison with VRC01 class antibodies, IOMA may overcome the N276-glycan barrier more readily: CDR L1 in IOMA (and in most other antibodies from VL2-23 and related germline genes) has a short α-helix that can accommodate the N276 glycan with fewer mutations than required by VRC01 class antibodies (47). In terms of features most favorable to epitope-based vaccine design, we note that antibody VRC34.01 had few epitope segments, low epitope-glycan content, and high epitope-conformational variability (48). These features may suggest that vaccine design based on the epitope of antibody VRC34.01 may be especially promising. Indeed, we recently achieved the elicitation of antibodies in mice, guinea pigs and rhesus macaques that neutralized diverse HIV-1 Env isolates through immunogen design based on the epitope of VRC34.01 (49). Thus, in addition to revealing features of paratope and epitope that allow for immune recognition of the prefusion closed spike, the structural survey also provides insight into which antibody templates are most suitable for vaccine design.

## Materials and Methods

### Structural Dataset and Selection of Structural Representative for Each of the 20 Unique HIV-1 Antibody Classes

All the antibody-antigen complex structures were collected from Protein Data Bank (PDB) (50) using the key words “HIV-1 antibody” on 28 February 2018. Structures that contain antibody alone were excluded. Non-human and synthetic antibodies were also excluded. The final comprehensive list of current deposited HIV-1 human antibody-antigen complex structures is provided in **Dataset S1**. Only the antibodies that target envelope trimer neutralize Tier 2 HIV-1 isolates were considered. To focus on broadly neutralizing HIV-1 antibodies, a neutralizing breadth cutoff of >25% was also used. Based on ontogenies in separate donors, we defined 20 unique classes that recognize the prefusion-closed Env trimer (**Fig. 1A**). For each antibody class, we selected the most informative structure based on the following criteria: 1) Resolution cutoff for X-ray crystal structures < 3.9 Å and < 4.5 Å for cryo-EM structures, allowing for a margin of 0.5 Å in resolution if a high resolution antibody-epitope complex were also available. 2) The complex structure of the trimer with highest resolution was selected, followed by glycosylated monomer, by deglycosylated monomer, or by scaffolded domain with highest resolution. The final PDB IDs we selected included 5V8L (18) (for PGT145), 3U2S (7) (for PG9), 5VGJ (34) (for VRC38.01), 5CEZ (51) (for 3H+109L and 35O22), 5ACO (52) (for PGT128), 4JM2 (53) (for PGT135), 1OP5 (19) (for 2G12), 4YE4 (for HJ16), 5T3Z (47) (for IOMA), 5F9O (25) (for CH235.09), 5FYJ (10) (for VRC01), 4YDJ (32) (for VRC13.01), 5VN8 (54) (for b12), 4JAN (28) (for CH103), (32) 4YDK (for VRC16.01), 6BF4 (11) (for VRC-PG05), 5FUU (55) (for PGT151), 5I8H (22) (for VRC34.01), and 5CJX (56) (for 8ANC195).

### Epitope and Paratope Buried Surface Area Calculations

The buried surface area between antibody and antigen was calculated using program NACCESS (57). The epitope residues and paratope residues for each antibody were defined as residues with non-zero buried surface area. For 2G12, the epitope residues were defined as glycans N295, N332, and N392, based on (20).

### Structural Definition of HIV-1 Env Broadly Neutralizing Antibody Categories

The structural representative of the 20 unique antibodies were clustered using hierarchical clustering, with the distance between each of the antibody is defined as 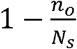, where *n_o_* is the number of overlapping epitope residues, and *N_S_* is the number of epitope residues for the antibody with the least number of epitope residues of the two.

### Display of Epitope Residues on Prefusion-Closed HIV-1 Env

The BG505 SOSIP trimer structure (PDB:5FYL) was used as the template with each glycan (with the addition of glycans 241 and 332) modeled as MAN5. Amino acid and glycan epitope residues were colored based on the structurally defined antibody category. Epitope residues with more than 5 Å C-alpha deviation from PDB:5FYL were excluded. For residue positions shared in epitopes from multiple antibody categories, the color of the antibody category with the antibody that has the largest epitope surface area was used.

### Antibody Expression and Purification

The 20 representative antibody heavy and light chain expression constructs were synthesized (Gene Universal Inc., Newark, DE) and cloned into pVRC8400 expressing vector. For antibody protein expression, 1.5 ml of Turbo293 transfection reagent (Speed BioSystems) were mixed into 25 ml Opti-MEM medium (Life Technology) and incubated for 5 min at room temperature. 500 μg of plasmid DNAs (250 heavy chain and 250 μg of light chain) were mixed into 25 ml of Opti-MEM medium in another tube. Then the diluted Turbo293 were added into Opti-MEM medium containing plasmid DNAs. Transfection reagent and DNA mixture was incubated for 15 min at room temperature, and added to 400 ml of Expi293 cells (Life Technology) at 2.5 million cells/ml. The transfected cells were cultured in shaker incubator at 120 rpm, 37 °C, 9% CO2 overnight. On the next day of transfection, 40 ml of AbBooster medium (ABI scientific) were added to each flask of transfected cells and the flasks were transferred to shaker incubators at 120 rpm, 33 °C, 9% CO2 for additional 5 days. At 6 days post transfection, antibodies in clarified supernatants were purified over 3 ml Protein A (GE Health Science) resin in columns. Antibody was eluted with IgG elution buffer (Pierce), immediately neutralized with one tenth volume of 1M Tris-HCL pH 8.0. The antibodies were then buffer exchanged in PBS by dialysis, adjusted concentration to 1.0 mg/ml and filtered (0.22 μm) for neutralization assays (58) (45).

### Virus Neutralization

Single-round-of-replication Env pseudoviruses were prepared, titers were determined, and the pseudoviruses were used to infect TZM-bl target cells as described previously in an optimized and qualified automated 384-well format (59). Briefly, antibodies were serially diluted, a constant amount of pseudovirus added, and plates incubated for 60 minutes; followed by addition of TZMbl cells which express luciferase upon viral infection. The plates were incubated for 48 hours and then lysed, and luciferase activity measured. Percent neutralization was determined by the equation: (virus only)-(virus+antibody)/(virus only) multiplied by 100. Data are expressed as the antibody concentration required to achieve 50% neutralization (IC50) and calculated using a dose-response curve fit with a 5-parameter nonlinear function. We used a previously described panel (5, 34, 60) of 208 geographically and genetically diverse Env pseudoviruses representing the major subtypes and circulating recombinant forms.

The IC_50_ values reported here are from the complete set of 208 viruses run at the VRC. In some cases, multiple runs were averaged. The values used here differed slightly, generally within 2 fold of previous median IC_50_, from previously reported values in references (5, 7, 11, 22, 24-40). The differences have two sources: in earlier publications, the panels contained between 178 and 198 of the 208 viruses; and because the neutralization IC_50_ values are known to vary up to 3-fold between repeat assays, as documented in (59).

### CDR H3 Length, Germline Gene Assignment, N-Insertion, and SHM Calculations

Antibody CDR H3 length, germline gene assignment, N-insertion, and SHM Calculation was performed using the IMGT/HighV-QUEST webserver (http://www.imgt.org/HighV-QUEST/).

### Number of Protruding Loops

CDR loops (i.e. CDR L1, CDR L2, CDR L3, CDR H1, CDR H2, CDR H3) were inspected manually, and two criteria were used in defining whether a CDR loop would be considered as protruding: 1) the loop extends at least 5Å beyond a plane defined by the average extent of the six antibody CDRs, and 2) the loop contacts with HIV epitope residues.

### Epitope Conservation Calculations

The conservation of each HIV-1 Env residue was calculated using the entropy function of the R package bio3d (H.norm column), with the addition of glycan as residue type. The calculation was based on the filtered web alignment of the year 2016 from the Los Alamos HIV sequence database (http://www.hiv.lanl.gov/). The epitope conservation was defined as the average conservation for each residue, weighted by epitope surface area. The average Env surface conservation was defined as the average conservation for each Env residue, weighted by accessible surface area as determined by NACCESS (57).

### Average Glycan Surface Calculations

The glycan surface area of the HIV-1 Env prefusion-closed trimer was calculated applying a two-step procedure. First, we approximated the accessible surface area (ASA), using NACCESS program (57) with the default radius of 1.4 Å, for all protein as well as glycan residues based on a molecular dynamics simulation with a 500-nanosecond timescale (10). Second, we determined the percentage of the glycan surface area, by dividing the ASA of glycan residues by the sum of the ASAs of protein and glycan residues.

### Number of Epitope Segment Calculations

The number of epitope segment is defined as number of continuous stretches on epitope residues, allowing skipping of one to two residues, which in total have an epitope surface area of greater than the specified threshold (25Å^2^ or 100 Å^2^).

### Epitope Structural Variation Calculations

The epitope structural variation for each antibody class was determined by the averaged root-mean-square-deviation of recognized epitope amino acid residues computed pairwise among category members and the unliganded structure (PDB ID: 4ZMJ) (33) using PyMOL software. Antibody 2G12 conformational variability was not calculated due to lack of amino acid epitope residues.

### Correlation Analyses

Pearson correlation for every pair of antibody properties listed in **Fig. 2** and **3** were calculated. b12 class was not included in the correlation calculation as b12 was derived from phage display. For correlations involving total epitope surface, paratope surface, and number of epitope segment calculations, PG9, VRC38.01, PGT135, H16, CH235, VRC13.01, CH103, VRC16.01, and VRC-PG05 were excluded as their representative structures were determined with partial Env domain. For correlations involving epitope glycan composition, HJ16, CH235, VRC13.01, CH103, and VRC16.01 were excluded as their representative structures were determined with deglycosylated gp120 monomers.

### Non-HIV-1 Antibody Sequence Dataset and V-gene Frequency Estimation

Heavy and light chain NGS samples (43) from 454 pyrosequencing downloaded from Short Read Archive and processed to calculate average SHM (**Table S1A – S1C**) and HV germline frequency (**Table S1D**). The next-generation sequencing (NGS) data from healthy donor performed with 5′ primer-adapters should avoid bias of germline gene usage. All heavy chain reads shorter than 350 nucleotides were removed, and all lambda and kappa chain reads longer than 300 nucleotides were kept. Germline genes were assigned to all filtered reads using IgBLAST (61). After assigned all reads, an in-house python script applied to process IgBLAST output, and non-IG sequences were removed. The variation between alleles were ignored, and we treated those alleles as identical VH gene. The VH gene frequency was calculated by the number of reads of VH germline divided by the total good IG sequences.

### SHM Rarity

Rarity score of a somatic hypermutation was calculated as 1-frequency of the SHM observed in gene-specific substitution profiles (43). SHMs with rarity score lower than 0.5% were counted as rare SHMs.

### Non-HIV Antibody-Antigen Structural Dataset

A non-redundant non-HIV-1 Env antibody/antigen structural dataset was obtained from SAbDab (62), using the “Non-redundant search” function (Antibody identity = 90%, Antigen identity = 99%, for protein antigens, Resolution = 3.9 Å). The list was further filtered down by keeping only the human antibodies, and removing all HIV antibodies, resulting a total of 81 antibody/antigen complexes (**Dataset S2**).

### Non-HIV Glycoprotein Structural Dataset

To understand the difference of epitope glycan composition between HIV-1 antibodies and non-HIV-1 glycoprotein, a non-HIV-1 glycoprotein structural dataset was assembled with 16 antibodies complexed with one of the following glycoproteins: Ebola virus glycoprotein, Herpes simplex viruses −2 gD, Human Metapneumovirus fusion protein, Hepatitis C virus envelope glycoprotein E2 core, dengue virus type 2 envelope glycoprotein, Human cytomegalovirus glycoprotein B, Respiratory syncytial F, Human cytomegalovirus pentamer, and influenza hemagglutinin **(Dataset S5)**. To avoid the dataset over-represented by influenza hemagglutinin, only four structures (the number of structures for the next abundant antigen) were selected. Resolution cutoff for X-ray crystal structures < 3.9 Å and < 4.5 Å for cryo-EM structures.

## Acknowledgements

We thank J. Stuckey for assistance with figures, and members of the Structural Biology Section and Structural Bioinformatics Core, Vaccine Research Center, for discussions or comments on the manuscript. We thank H. Mouquet and M. Nussenzweig for the nucleotide sequences of antibody 10-1074. We also thank J. Baalwa, D. Ellenberger, F. Gao, B. Hahn, K. Hong, J. Kim, F. McCutchan, D. Montefiori, L. Morris, J. Overbaugh, E. Sanders-Buell, G. Shaw, R. Swanstrom, M. Thomson, S. Tovanabutra, C. Williamson, and L. Zhang for contributing the HIV-1 envelope plasmids used in our neutralization panel. Support for this work was provided by the Intramural Research Program of the Vaccine Research Center, National Institute of Allergy and Infectious Diseases, and by IAVI Neutralizing Antibody Consortium (NAC).

## Author contributions

G.-Y.C., L.S. and P.D.K. designed research; G.-Y. C., J.Z., R.R., C.-H. S., Z. S., A. P. W., B.Z., T.Z., R.T.B., N.A.D. and .M.L. performed research; G.-Y.C., L.S., and P.D.K. wrote the paper, with all authors providing comments or revisions.

